# Disturbances can facilitate prior invasions more than subsequent invasions in microbial communities

**DOI:** 10.1101/2023.12.05.569871

**Authors:** Luke Lear, Elze Hesse, Angus Buckling

**Affiliations:** Environment and Sustainability Institute, University of Exeter, Penryn, Cornwall, United Kingdom, TR10 9FE

**Keywords:** invasion, disturbance, order, timing, propagule pressure, microbial community, multiple-stressors

## Abstract

Invasions by microorganisms are commonly found to benefit from disturbance events. However, the importance of the relative timing and order of the invasion and disturbance for invader success remains uncertain. Here, by invading a stably coexisting community of five bacterial species at four different propagule pressures on eight separate occasions – four pre-disturbance and four post-disturbance – we experimentally tested the importance of timing and order for the invader’s success. Furthermore, we quantified the impact of the invader on the composition of the resident community. Across a range of invader densities, both pre- and post-disturbance invader success was greatest the closer in time to the disturbance. While the greatest invasion success occurred when the invasion immediately followed the disturbance, invasion resistance rapidly recovered, such that the three subsequent invasions had negligible success. As a consequence, overall invader success was greatest pre-than post-disturbance. We found that invasion day as well as success significantly affected resident diversity, such that it was lowest in communities invaded immediately after the disturbance, but was overall lower in communities invaded pre-disturbance. Overall, these results demonstrate that invading immediately after a disturbance is highly beneficial for invader success and results in the biggest change to community composition. Importantly however, they also show that this window of opportunity is very brief, and that, on average, an invader will have a greater success and effect on community composition if it invades before a disturbance.

## Introduction

Invasions of microbial communities are ubiquitous events with potentially significant consequences for the diversity and function of those communities (1, 2). Furthermore, microbial invaders can be beneficial biocontrol agents (3, 4), as well as play an important role in both disease acquisition (invasion by a pathogen) (5, 6) and prevention and recovery (the use of probiotics) (7-9). As a result, understanding how and when novel microbes may successfully invade a community is an important goal. Moreover, the control and replication afforded by microbial communities make them useful systems for providing general insight into ecological processes.

One factor that has frequently been shown to positively affect invader success is disturbance – events that destroy biomass and consequently change the availability of resources and habitat (10-13). It is the associated changes in resource abundance, the opening up of habitats, and the reduction of both resident priority and dominance effects that are the proposed key mechanisms for how disturbances can facilitate invasions (12-15). These mechanisms reduce the resident community’s invasion resistance by reducing competition between the invaders and resident community members (11, 13, 16).

Whilst previous work has shown that invaders are often successful immediately after a disturbance (17-19), it is uncertain how either the relative timing or the order of an invasion and a disturbance event impact the invader’s success. For example, it is unclear whether disturbances can facilitate invasions by exotic species already present in the community, how long invasion resistance takes to recover, and whether it recovers linearly with time since disturbance or if there is only a narrow window of reduced resistance. Understanding these aspects of microbial invasions is crucial for determining how long disturbed communities are vulnerable for, and how consistent this vulnerability is through time. Moreover, it will address whether conclusions resulting from previous experimental invasion work, of which the majority focuses on disturbance and invasion events occurring simultaneously (17-19), are supported when the disturbance and invasion events are asynchronous. Furthermore, it is important that we understand if the interactive effect of invasion and disturbance on resident biodiversity depends upon their relative order and timing, as both stressors are increasing as a consequence of anthropogenic activities (20-22).

A crucial factor for invader success that may influence the importance of the relative invasion timing is propagule pressure – the number of invaders entering the novel environment (23-26). This is because smaller invading populations, which are less likely to be genetically diverse than larger populations, are more likely to have both reduced establishment probability and reduced long term invader success (25, 27, 28). Furthermore, smaller invading populations are more likely to suffer from Allee effects, such as being stochastically removed by either demographic or environmental causes, including the disturbance event (24, 29, 30). Therefore, it is probable that having a large invading population is crucial for surviving to, and through, the disturbance, and that post-invasion disturbances may be detrimental for the success of small invading populations.

Here, we take an experimental approach to test the importance of the timing and order of an invasion event in relation to a disturbance for the success of the invader, and whether this changes with invader density. We additionally determine if the timing of invasions affects resident community composition over and above invasion success. We did this by invading a resident community of five stably coexisting bacterial species (31) on one of seven days and at four different invader densities (propagule pressures). On the fourth invasion day, we disturbed all communities, and carried out an invasion event both before and after the disturbance. At the end of the experiment we measured the density of the invader and the composition of the resident community.

## Methods

### Bacterial strains

We used the same model resident community and invader strain as in previous work (32). The resident community used is comprised of five bacterial species previously isolated from the same compost sample: *Achromobacter sp*., *Ochrobactrum sp*., *Pseudomonas sp*., *Stenotrophomas sp*., and *Variovorax sp*.. These species have been shown to stably coexist in diluted tryptic soy broth (TSB) media and, crucially, to have visually distinct colony morphotypes when plated onto King’s medium B (KB) agar (31). Monocultures of each of the five species were grown for 48 hours at 28°C in shaking (180rpm) glass vials containing 6mL of diluted (1/64^th^) TSB. Using optical density measurements at 600nm, the monocultures were diluted to 10^5^ colony forming units (CFU) per mL and combined to form a mixture of each of the five species. Next, 60μL of the mixture was added to 192 microcosms containing 6mL of diluted TSB; these were kept static throughout the experiment with loose lids to allow oxygen transfer. A further 42 microcosms were setup; on each invasion day 6 of these were destructively sampled in order to quantify an approximate value of resident density at the time of invasion. This allowed us to test whether the initial invader proportion varied between the day treatments at a given propagule pressure.

We used *Pseudomonas aeruginosa* PA01 as our invader (33). *P. aeruginosa* can successfully invade a range of environments, and has previously been found to successfully invade our model resident community under some disturbance regimes, but not under others (32). In the experimental conditions used here, this strain has a growth rate around three times that of the whole resident community (supplementary information, section 1). In order to easily distinguish this strain from the resident community it is lacZ-marked: this causes a blue colour change when plated on agar containing X-gal (33). Monocultures of *P. aeruginosa* were grown in shaking glass microcosms containing 6 mL of undiluted TSB for 24 hours at 28°C. We grew the invader in undiluted TSB as we wanted to achieve a high invader density in order to be able to have a range of propagule pressures. However, as the addition of resources alongside the invader would in itself be equivalent to a disturbance that may influence invader success and/or the composition of the resident community (12), we ‘washed’ the invader before addition to remove any growth medium. To do this, the monocultures were fully homogenised, before 2mL was centrifuged at 14800rpm for 2 minutes, the supernatant was then removed and the pellet re-suspended in 2 mL of M9 salt buffer. The density of this inoculate was then standardised to an optical density of 0.6 ± 0.07 at 600nm, which resulted in mean CFU measurements of 6.41 × 10^7^ ± 2.96 × 10^7^ SD after 48 hours incubation on King’s medium B (KB) agar at 28°C. To achieve a range of propagule pressures, 60 μL of the washed invader was added per invasion event either undiluted (high invader density), diluted 10-fold (medium-high invader density), diluted 100-fold (medium-low invader density) or diluted 1000-fold (low invader density).

### Experimental design

A total of 32 treatments were used, each replicated 6 times for a total of 192 microcosms. This included 8 separate invasion events which occurred on either day 5, 6, 7, 8 (either before or after the disturbance), 9, 10 or 11, at the four propagule pressures. All microcosms were disturbed on day 8; this consisted of fully homogenising each microcosm, before 60 uL (1% by volume) was transferred into fresh media. Disturbing in this way represents a non-specific pulse disturbance event that removes 99% of the community regardless of its density (34). In addition to removing any growth media, we limited any physical disturbance during invasion events (e.g. to any biofilms) by pipetting the invader down the side of the microcosm and not homogenising the culture before or after (12).

On day 16, all microcosms were thoroughly homogenised, mixed with glycerol to a final concentration of 25%, and frozen at -70°C. Thawed samples were diluted in M9 salt buffer to a dilution of 10^−5^, plated on KB agar containing X-gal and incubated for 48 hours at 28°C, before the number of CFUs was quantified. We note that one replicate from the day 8 post-disturbance, medium-high invader density treatment was lost during the experiment.

### Statistical analysis

We first determined if the resident density differed between invader day treatments (i.e. if the resident population density was statistically similar at the time of invasion across invader days). To do this we used a linear model, with resident population density (number of CFU) as the response variable and day (factor) as the explanatory variable.

Invader success was quantified as final invader abundance. This metric is common among the invasion literature, and has been used for both plant (35) and microbe studies (27). We chose this metric over those using a measure of invader fitness or growth rate, as used in some previous studies (12, 34, 36), because the differences in initial invader density (i.e. due to the propagule pressure treatments), combined with the maximum value of invader (e.g. its carrying capacity or becoming 100% of the community through displacing the residents) being equal across treatments, results in the maximum possible fitness or growth rate values being different between these treatments. For example, invaders going from 0.001% to 100% and 1% to 100% of the community will have very different fitness values, but it is difficult to know how meaningful that difference is, as the fitness of the latter cannot be as large as the former, and so is likely to be underestimated. As the final invader abundance had a high number of zeros and the variance in final invader abundance (894.1) was greater than its mean (18.18), we analysed success using a negative binomial generalised linear model with invasion day (factor), propagule pressure (factor), and their interaction as the explanatory variables. We additionally used a generalised linear mixed model using Template Model Builder with invader and resident density as a binomial response, and day as a dispersion parameter (due to zero inflation), to test invasion day and propagule pressure effects on the proportion of the invader in the community.

To test the effect of invasion day and propagule pressure on the biodiversity of the resident community, we first calculated diversity using the Simpson’s index 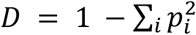 where *p*_*i*_ is the proportion of the *i*^*th*^ species of the resident community (37). We then used a linear model with invasion day (factor), propagule pressure (factor), and their interaction as the explanatory variables. Finally, to test whether treatment effects on diversity were due to treatments affecting invader success, we added invader abundance as a covariate in our model of diversity.

All analysis were done in R version 4.1.0 (38). We used the ‘*MASS*’ package (39) for the negative binomial generalised linear model, the ‘*glmmTMB*’ package (40) for the generalised linear mixed model using Template Model Builder, and the ‘*DHARMa*’ package to check the residual behaviour of our models. We sequentially removed model terms and compared model fits using either F or χ^2^ tests in order to arrive at the most parsimonious model. We then used the ‘*emmeans*’ package (41) to calculate contrasts, with p values adjusted for multiple testing using the Bonferroni method where appropriate. All reported CFU count data is the number of CFU present on the KB agar plate; information on how to convert this to CFU/mL is in the associated code (42).

## Results

### Invaders added pre-disturbance were on average more successful and benefited more from greater propagule pressure

Invader success was significantly different between invasion days (day main effect on invader density: *χ*^2^ = 683.8, d.f. = 7, p < 0.001), with pre-disturbance invasions having on average much greater success than post-disturbance invasions (mean ±SD invader abundance when invading on days 5, 6, or 7 = 14.4 ± 13.8 CFU; mean ±SD invader abundance when invading on days 9, 10, or 11 = 1.17 ± 1.22 CFU). An exception to this was when invasions occurred immediately after the disturbance on day 8, as this treatment had the highest average success (mean CFU ±SD: 69.4 ± 45.2), followed by the invasion event that occurred immediately before the disturbance (mean CFU ± SD: 31.3 ± 29.0; (Fig.1).

As expected, increasing propagule pressure had the overall effect of significantly increasing invader success (propagule pressure main effect: *χ*^2^ = 150.4, d.f. = 3, p < 0.001). However, we found propagule pressure and invasion day to significantly interact (day-propagule pressure interaction: *χ*^2^ = 81.72, d.f. = 21, p < 0.001; Fig.1). In this interaction, we found that in general increasing propagule pressure had a much greater effect on invader success on pre-disturbance days than on post-disturbance days, with the invasion event immediately after the disturbance once again being an exception. Pre-disturbance, final invader abundance significantly increased between at least two propagule pressure treatments on each day, with the difference in success between the high and low, and high and medium-low invader density treatments always being significant (p=<0.005 for all contrasts). Success only differed between high and medium-high invader density on days 7 (p=0.0008) and 5 (p=0.021). Post-disturbance, propagule pressure was only significantly different between high and medium-low invader density on day 9 (p=0.0098; p=>0.051 for all other contrasts). For the invasion event immediately after the disturbance, we note than an outlier value of 158 invader CFU in the low propagule pressure treatment (mean without outlier = 15.2) altered the comparisons between the treatments. With the outlier present, there was no statistical difference between any treatments (p=>0.051 for all contrasts), with it removed however, the low invader density treatment had significantly lower success than the medium-low (p=0.0017), medium-high (p<0.0001), and high (p<0.0001) invader density treatments; all other contrasts were not significant (p=>0.081 for all contrasts). Consequently, our results show that propagule pressure is not always a good predictor of invader success, with the success of at least 25% of the invasion events (days) here not differing as a function of propagule pressure. Notably, those invasion events where propagule pressure did not influence success were all post-disturbance. Across all propagule pressures, the invasion event immediately after the disturbance was the only one to have 100% establishment success (a final invader abundance of at least 1), with the three subsequent events having an average establishment success of <50%, and the four pre-disturbance events having an average establishment success of >80% (Fig. 1).

**Figure 1.**
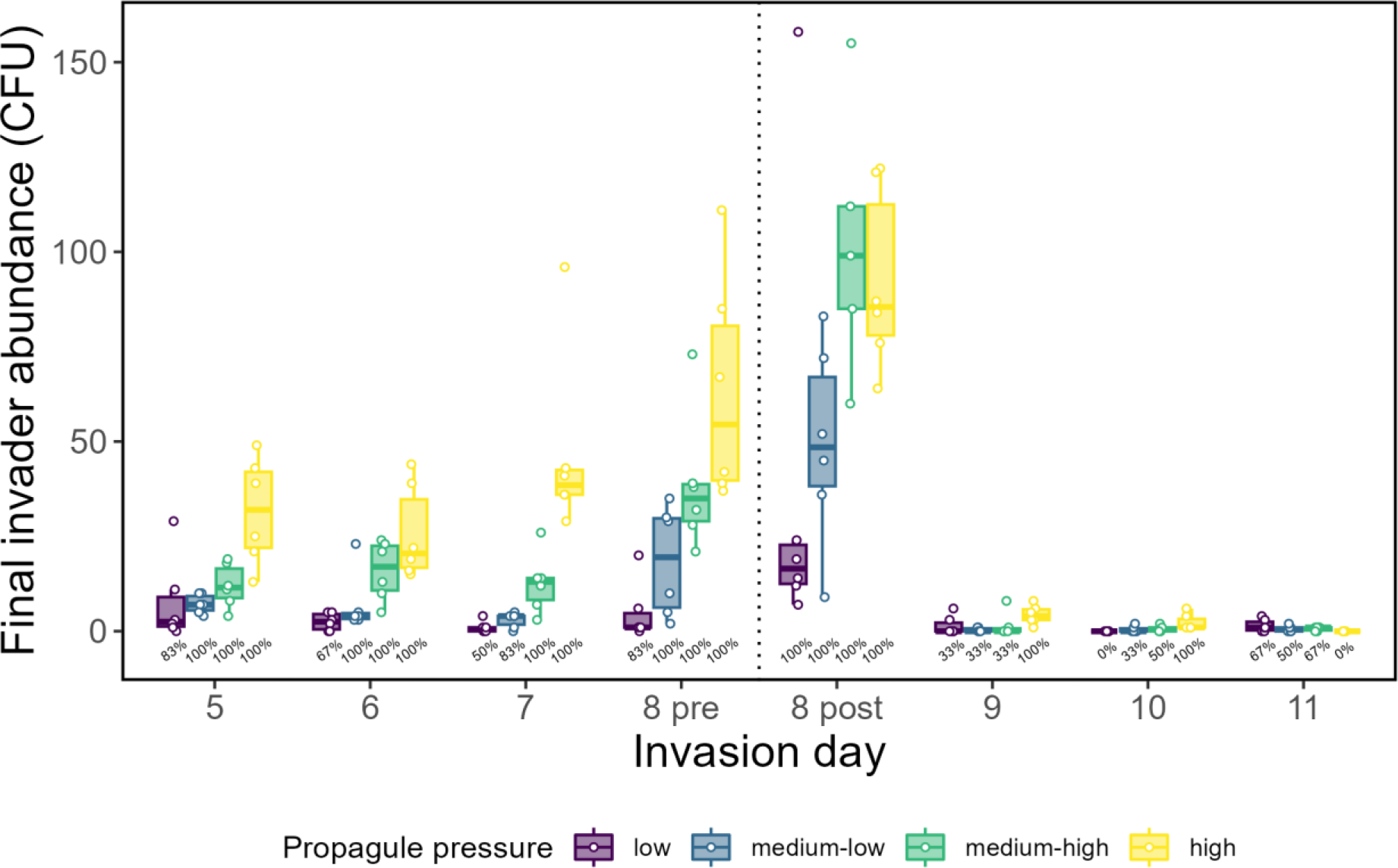
The number of invader colony forming units (CFU per agar plate) on day 16 in communities that have been invaded on different days relative to a pulse disturbance event on day 8 (represented by the vertical dotted line). Colours represent different propagule pressure treatments: purple = low invader density, blue = medium-low invader density, green medium-high invader density and yellow = high invader density. Circles show individual replicates (n = 6) and numbers show establishment success (percentage of replicates with at least 1 final invader present): 0% = no invader present in any of the 6 replicates; 100% = invader present in all 6 replicates.

Repeating these analyses with proportion of invader in the community instead of final invader abundance as the dependent variable, we found the same trend to occur (day-propagule pressure interaction: *χ*^2^ = 44.89, d.f. = 21, p < 0.002; supplementary information Fig.2). These results show that the timing of an invasion event relative to a disturbance is a crucial determinant of both its direct success, and the effect of propagule pressure. Moreover, they demonstrate that disturbances can be highly beneficial for invaders, even if they occur multiple generations after the invasion event, and that although the post-disturbance window of opportunity offers very high success, it is very brief.

We note that the density of the resident communities (mean total CFU per vial ± SD: 1.46 × 10^8^ ± 4.51 × 10^7^) did not significantly differ over the seven days the invaders were added (F_6,35_=1.25, p = 0.30), and consequently any differences in invader success across days within each propagule pressure treatment are not due to variation in the initial proportion of the invader in the community.

### Resident diversity decreased with invader abundance

Both the day the invader was added and propagule pressure had significant main effects on resident diversity (day main effect: F_7,159_=23.14, p <0.001; propagule pressure main effect: F_3,159_=11.94, p < 0.001; Fig. 2). However, they significantly interacted in a way that mirrored treatment effects on invader success (day-propagule pressure interaction: F_21,159_=2.83, p = 0.001), such that increasing propagule pressure generally decreased diversity on days 5, 6, 7, and 8, but this propagule effect disappeared on days 9, 10 or 11. When averaging over the effect of propagule pressure, we found that the communities invaded immediately after the disturbance had significantly lower diversity (mean Simpson’s index ± SD = 0.36 ± 0.19) than all other invasion events (p=<0.001 for all contrasts). Moreover, we also found that the community invaded immediately before the disturbance had a significantly lower diversity (mean Simpson’s index ± SD = 0.50 ± 0.18) than all other communities (p=<0.0034 for all contrasts) except communities invaded on day 7 (mean Simpson’s index ± SD = 0.56 ± 0.16; p=0.082), and that communities invaded on day 7 had lower diversity than those invaded on day 9 (mean Simpson’s index ± SD = 0.64 ± 0.07; p=0.0174). However, communities invaded on day 6 did not significantly differ in diversity to those invaded on day 10 (p=0.192), nor did communities invaded on day 5 to communities invaded on day 11 (p=0.079).

**Figure 2.**
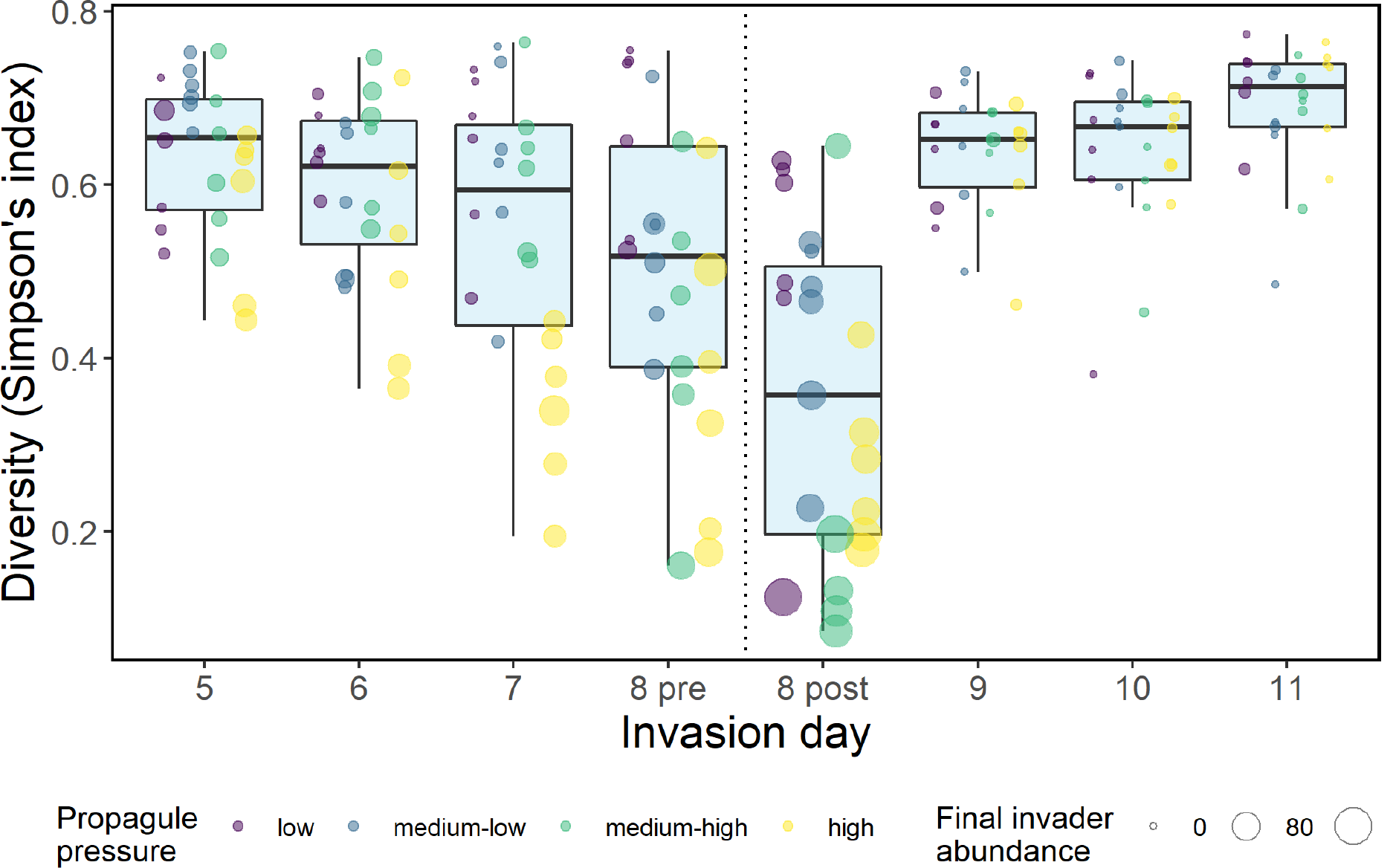
The resident diversity (Simpson’s index) of communities on day 16 that have been invaded on different days by invading populations at different densities, and exposed to a disturbance on day 8 (represented by the vertical dashed line). Colours represent different propagule pressure treatments: purple = low invader density, blue = medium-low invader density, green = medium-high invader density and yellow = high invader density; circles show individual replicates (n = 6), and their size corresponds to final invader abundance.

To determine if diversity decreased as a direct consequence of the treatments, or purely as an indirect effect due to their effect on invasion success, we added invader abundance as a covariate in our model of diversity. We found propagule pressure to no longer be a significant predictor of diversity (F_3,179_=0.54, p=0.066). However, both invasion day and invader abundance significantly affected diversity (day: F_7,182_=28.96, p<0.001; invader abundance: F_1,182_=136.20, p<0.001), with, on average, the Simpson’s index of a community being reduced by 0.004 for every additional final invader CFU present. Consequently, communities invaded immediately after a disturbance were the least diverse, but in general communities invaded post-disturbance were more diverse than those invaded pre-disturbance, especially at high propagule pressure. These results demonstrate that the same invader can have different effects on community diversity, depending on the timing of the invasion event relative to a disturbance.

### Residents were most likely to become extinct when the invasion and disturbance occurred on the same day

As we found diversity to be very low (values of Simpson’s index <0.02) in multiple final communities, we further explored changes in the resident composition by quantifying the number of times a resident species became extinct (had 0 CFU) in a community. A total of 64 resident extinctions occurred across all 32 invaded treatments (6.67% of species; Fig. 3). The majority of these extinctions (n=49 or 76.6%) occurred when the invasion and disturbance event occurred on the same day, with 11 happening when the invader was added before the disturbance and 38 happening when it was added after. Grouping the remaining six invasion events into either invaded pre-disturbance or post-disturbance, we found that twice as many extinctions occurred when the invader was added before the disturbance (n=10) than when it was added after (n=5). We found the resident species to be affected differently: *Stenotrophomas* did not become extinct in any community; *Pseudomonas* was severely affected by invasion immediately before the disturbance, becoming extinct in all communities invaded with the highest propagule pressure in this treatment; and *Ochrobactrum, Achromobacter*, and *Variovorax* were most vulnerable to extinction when the invasion immediately followed the disturbance. These results show that the timing and order of an invasion and disturbance event can have major and long-term implications for the composition of the resident community by causing multiple species to become extinct.

**Figure 3.**
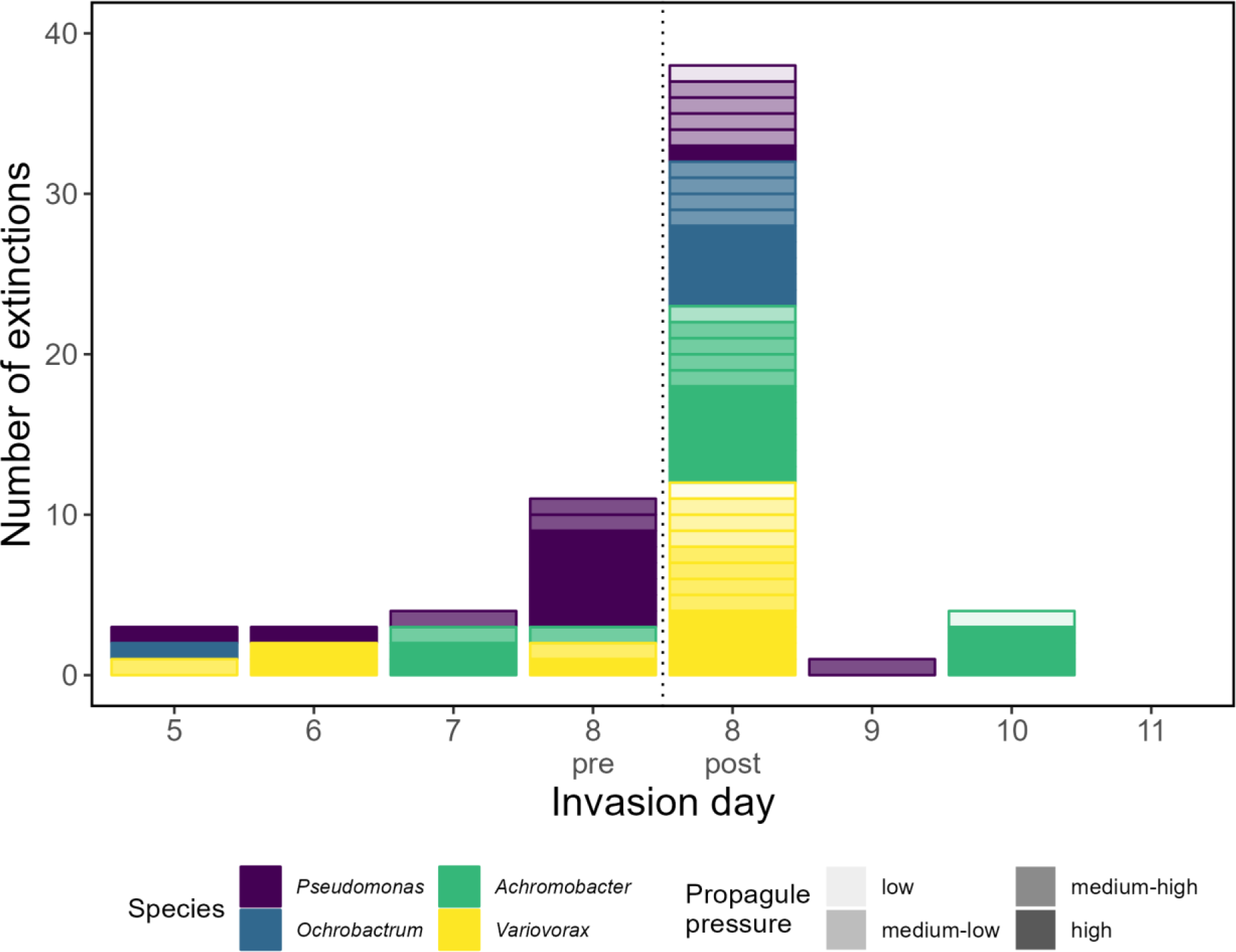
The number of resident extinctions in disturbed communities of five bacterial species that have been invaded on different days. Colours show the different species: yellow = *Variovorax*, green = *Achromobacter*, blue = *Ochrobactrum*, and purple = *Pseudomonas*. We note that *Stenotrophomonas* did not become extinct in any treatment. Opacity shows the density of the invader added, with less transparent colours showing greater propagule pressure. The vertical dashed line shows when the disturbance took place (day 8).

## Discussion

Here, we experimentally tested how the relative timing and order of an invasion in relation to a disturbance event affects the success of the invader, and whether this changes with propagule pressure. Specifically, we tested if pre-disturbance invasion events can benefit from the disturbance, and how quickly post-disturbance invasion resistance is restored. We found invasion day to be a significant predictor of invader success, with invaders added on the days prior to a disturbance having success on average twelve times greater than those added on post-disturbance days. However, invaders added on the day of the disturbance were exceptions to this, with those added immediately before the disturbance having an average success two times greater than all prior invasions, but only having an average success half that of the populations added immediately after the disturbance. Moreover, we found invasion day to significantly interact with propagule pressure, such that greater propagule pressure increased success on the days before the disturbance and immediately after it, but had very little effect on the subsequent post-disturbance days.

Surprisingly, we found the invasion event immediately before the disturbance to have greater average success than all other invasion events except the one that immediately followed the disturbance. This suggests that the benefit of disturbances to the fast growing invader used here outweighs the 99% reduction in their population caused by the disturbance. One plausible reason for this event having higher success than the previous invasion events is that without the benefit of disturbance, invader populations are outcompeted by the residents through priority and dominance effects that cause their populations to begin to decline (43, 44). However, when added immediately before the disturbance, they have yet to be excluded, and so remain at relatively high densities compared to the invaders added on previous days. These findings therefore suggest that if an invader can survive to and through a disturbance event, it is likely to benefit from the reduction in invasion resistance caused by that disturbance. However, by this time its population density has likely been reduced, and so populations invading immediately after the disturbance have greater success, especially as the latter will also be invading communities with reduced density. We note that in addition to the invader surviving the disturbance, it is important that their offspring do too. For example, trout invasions fail when flood disturbances kill their fry, but are successful if they breed after the flood season (45).

Our results show that the invasion resistance of a microbial community can recover rapidly, with very low invader success occurring when the invasion occurred just 24 hours after the disturbance. Moreover, this recovery was associated with a large reduction in the impact of the invader on the composition of the resident community. This finding may explain why higher frequencies of disturbance have been found to increase both invader success and impact on the community (32, 34, 46) – we show that a disturbance occurring is beneficial for an invader on the short term, but that the resident community can quickly restore its resistance. Therefore, as the disturbance frequency increases the number of times the invader benefits will also increase, and the period after the disturbance where success is low will decrease, as the time between disturbances is shorter.

Propagule pressure typically increases the success of an invader (*but see* (25, 47)), and this effect is typically much greater when going from low to medium invader density than from medium to high invader density (24, 25, 48, 49). Here, however, we demonstrate that the effect of increasing propagule pressure on invader success is not always consistent, and that it can depend on the relative timing of the invasion event. For example, consistent with previous work testing the interaction between disturbance and propagule pressure, we found propagule pressure to increase invader success when the invader was added on the same day as the disturbance (17-19). However, we found propagule pressure to not significantly affect the success of 25% of the invasion events, with the success of invasion events that occurred days after the disturbance not differing with invader density. We do show, however, that when propagule pressure does affect success, it is likely to plateau as the two highest propagule pressures only differed in success on one out of eight occasions. These findings could have important implications for the use of biological control agents, where a balance between high colonisation success and economic cost is crucial (50). For example, we demonstrate that large doses may result in negligibly more success than medium doses, and it is instead better to do two temporally separated medium introductions rather than one large introduction, as it increases the chance that the timing will be beneficial. This tactic may provide further advantage by restoring any lost genetic diversity in the initial invading population (51), or by reducing the probability of a chance event removing the whole invading population (50).

That the day the invader was added affected resident diversity over and above invader success, shows that timing is a crucial determinant of an invader’s impact on a resident community. This is demonstrated by the majority of resident extinctions occurring when the invasion and disturbance occurred on the same day. Consequently, our findings show that the impact of an invader on a community cannot always be predicted by its success, and that the vulnerability of a community to invader-induced diversity loss changes across a disturbance regime. The loss of resident diversity as a result of invasion supports previous findings that invaders are in themselves a disturbance (52-55). Therefore, our results provide further insight into the effect multiple disturbances may have on resident diversity, and demonstrate that the order, relative timing, and intensity can alter the direction and severity of their interaction for resident diversity. For example, we demonstrate that diversity is reduced the most when the invasion and disturbance occur on the same day. This suggests that the interaction between two disturbances on resident diversity may become more synergistic the closer they are temporally. Furthermore, our data show that the importance of the order of the disturbances decreases as they become more temporally separated. For example, we find that communities invaded immediately before and immediately after the disturbance have significantly different final diversity, as do communities invaded on day 7 compared to day 9; however, we find that communities invaded on day 6 do not have significantly different diversity to those invaded on day 10, nor those invaded on day 5 to day 10. Moreover, we show that the order and timing can alter the effect of increasing the intensity of the invasion disturbance (i.e. changes in propagule pressure) on diversity loss, as this effect is not consistent across the disturbance regime. This change in the degree of synergism between multiple disturbances for resident diversity with order, timing, and intensity may help to explain why there is not a consistent trend in how multiple disturbances interact (56).

Importantly, our results may have practical applications for the invasion of microbiomes. For example, our results demonstrate that disturbing a host microbiome, such as by taking antibiotics, could increase the risk of invasion by a pathogen, or cause a species already present to flourish and cause dysbiosis and/or further infection (57, 58). Therefore, we highlight a potential risk when using preventative antibiotic therapy (59, 60), and note our findings are likely to be more extreme when the invader is resistant to the disturbance (e.g. is resistant to the antibiotic used). Similarly, probiotic microorganisms are commonly consumed after antibiotic therapy to restore the health of the gut microbiome (61). However, our findings suggest that, assuming the probiotic has a degree of tolerance to the antibiotic, it may be more beneficial to consume them before taking the antibiotic, as the chance that they successfully establish is likely to be greater. Outside of host microbiomes, our results could enhance the function of industrial microbiomes. For example, the invasion of poor performing anaerobic digesters by communities from high performing digesters can lead to a beneficial increase in gas production (47); here we demonstrate that coinciding the introduction with a disturbance could lead to greater establishment, even at low propagule pressures.

In conclusion, our results show that the timing of an invasion event relative to a disturbance is crucial to both its success and impact on the resident community. We demonstrate that disturbances reduce community invasion resistance, resulting in the invaders arriving immediately after having the greatest success. However we also show that invasive species already present benefit from the reduced resistance. Consequently, it is important that efforts are made to cause minimal disturbance in recently invaded environments, and to consider that disturbing an environment may allow undetected invaders to flourish. This is particularly important when we consider that the beneficial window for invaders is open for much longer before the disturbance than after it.

## Data availability statement

All data and R code used to analyse it are deposited on Zenodo (42).

