## supplementary information for "Disturbances can facilitate prior invasions more than subsequent invasions in microbial communities"

Luke Lear[1]*, Elze Hesse[1] and Angus Buckling[1].

[1]Environment and Sustainability Institute, University of Exeter, Penryn, Cornwall, United Kingdom, TR10 9FE

This supplementary information contains information on the growth rates of the resident community used (all five species combined) and of the invader, *Pseudomonas aeruginosa*. We also provide details on the proportion of the invader in the final community. All associated data and code can be found on Zenodo (Lear *et al.* 2023).

### Section one – Growth rate analyses

#### *Methods*

We quantified the growth rates of six replicate populations of the invader and six replicates of 8-day old resident community set up as described in the *Bacterial strains* section of the main document. To do this, 20µL of each of the 12 cultures was transferred into separate wells of a 96-well plate containing 180µL of diluted (1/64$^{th}$) TSB media (three technical replicates per culture resulted in 36 wells being inoculated). The plate was then incubated in a Biotek spectrophotometer for 24 hours at 28°C with optical density readings taken at 600nm every ten minutes.

*Statistical analyses*

To analyse the growth rates of the invader and resident community, we first trimmed our dataset to leave the observations between one and twelve hours of growth, due to air bubbles inflating optical density readings outside of these times. We then subtracted the lowest value from each well from all other readings in that well, so that all curves started at the same optical density value (0). We then used the '*growthcurver*' package (Sprouffske & Wagner 2016) to calculate the growth rate, *r*, of the cultures, before averaging this value for the three technical replicates per culture. Finally, we used a linear model to test whether *r* was different between the invader populations and resident community.

*Results – The invader has a higher growth rate than the resident community*

We found a significant difference in the growth rate of the community and the invader ($F_{1,10}$=118.1, p<0.001; Fig. S1), with the invader growth rate (0.0264 ± 0.0041 SD) being over three times greater than that of the resident community (0.0079 ± 0.0008).

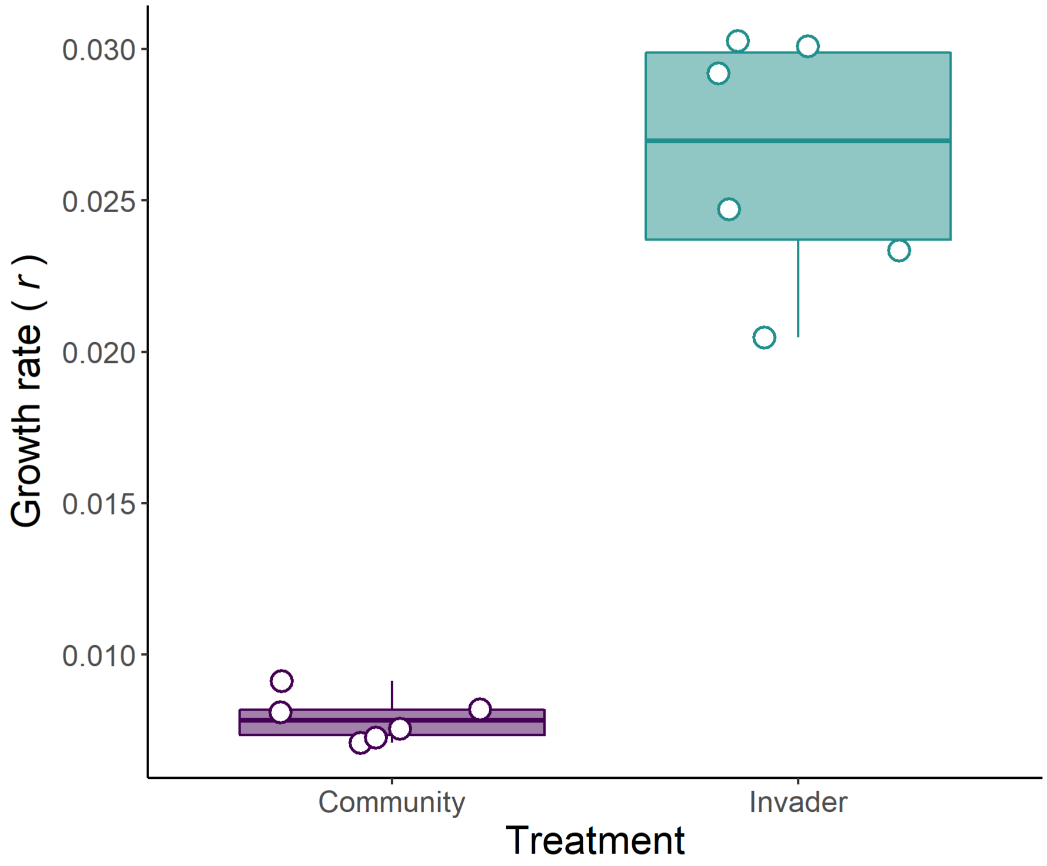

**Figure S1** The growth rate, *r*, of the resident community (purple) and the invader (green) quantified using optical density measurements. Circles show individual replicates (n = 6), and are an average of three technical replicates.

<u>Section two – final invader proportion</u>

To demonstrate that final invader abundance is a measure of invader success and not of some treatments being more productive than others (i.e. to confirm that the invader was more abundant relative to the residents), we repeated the invader success analysis using proportion of invader in the community (final invader CFU / (final invader CFU + final resident CFU)) instead of final invader abundance. Using a binomial generalised linear mixed model with invasion day, propagule pressure, and their interaction as dependent variables, we found the same trend to occur when using invader density (Fig. S2A) and proportion (Fig. S2B) (day-propagule pressure

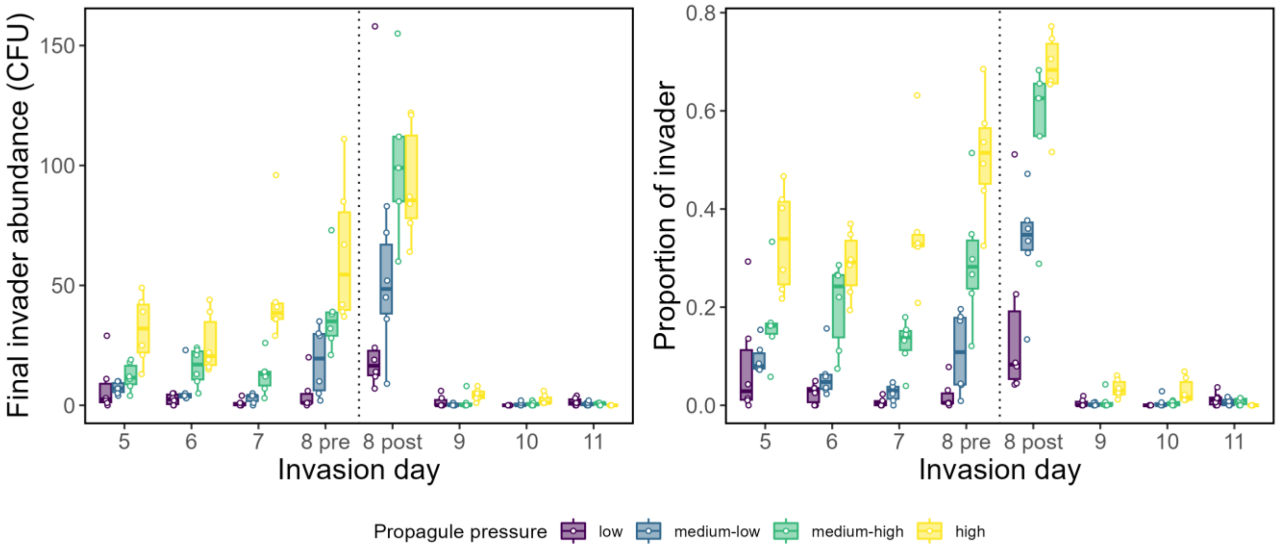


interaction: $\chi2 = 275.31$, d.f. = 21, $p < 0.001$).

**Figure S2** The success of an invader quantified on day 16 in communities that have been invaded on different days relative to a pulse disturbance event on day 8 (represented by the vertical dotted lines). Panel A shows the number of invader colony forming units (CFU), and panel B the proportion of invader in the community (final invader CFU / (final invader CFU + final resident CFU)). Colours represent different propagule pressure treatments: purple = low invader density, blue =

medium-low invader density, green medium-high invader density and yellow = high invader density. Circles show individual replicates (n = 6).
